# A systematic protocol to identify “clinical controls” for pediatric neuroimaging research from clinically acquired brain MRIs

**DOI:** 10.1101/2025.06.25.661530

**Authors:** Dabriel Zimmerman, Ayan S. Mandal, Benjamin Jung, Matthew J. Buczek, Jenna M. Schabdach, Shivaram Karandikar, Eren Kafadar, Margaret Gardner, Maryam Daniali, Laura Mercedes, Sepp Kohler, Leila Abdel-Qader, Raquel E. Gur, David Roalf, Theodore D. Satterthwaite, Remo Williams, Vivek Padmanabhan, Jakob Seidlitz, Lauren K. White, Susan Sotardi, J. Eric Schmitt, Arastoo Vossough, Aaron Alexander-Bloch

## Abstract

Progress at the intersection of artificial intelligence and pediatric neuroimaging necessitates large, heterogeneous datasets to generate robust and generalizable models. Retrospective analysis of clinical brain magnetic resonance imaging (MRI) scans offers a promising avenue to augment prospective research datasets, leveraging the extensive repositories of scans routinely acquired by hospital systems in the course of clinical care. Here, we present a systematic protocol for identifying “scans with limited imaging pathology” through machine-assisted manual review of radiology reports. The protocol employs a standardized grading scheme developed with expert neuroradiologists and implemented by non-clinician graders. Categorizing scans based on the presence or absence of significant pathology and image quality concerns facilitates the repurposing of clinical brain MRI data for brain research. Such an approach has the potential to harness vast clinical imaging archives – exemplified by over 250,000 brain MRIs at the Children’s Hospital of Philadelphia – to address demographic biases in research participation, to increase sample size, and to improve replicability in neurodevelopmental imaging research. Ultimately, this protocol aims to enable scalable, reliable identification of clinical control brain MRIs, supporting large-scale, generalizable neuroimaging studies of typical brain development and neurogenetic conditions. Studies using datasets generated from this protocol will be disseminated in peer-reviewed journals and at academic conferences.

## Introduction

A major challenge in the study of neurodevelopment is the need for large, diverse study samples to support valid scientific inference and generalizability. Recent work has argued that thousands of study participants are required for well-powered, replicable brain magnetic resonance imaging (MRI) studies in many contexts – given relatively small effect sizes observed in many well-powered studies (1). It is extremely difficult for individual research studies or even multi-site consortia to achieve such sample sizes. Though improved sampling schema hold the potential to partially mitigate this problem (2,3), brain MRI scans acquired for clinical purposes may provide a valuable alternative to augment sample sizes available for human brain imaging research. Despite logistical limitations of clinical brain MRIs (i.e., their lack of advanced multimodal sequences), valuable quantitative phenotypes can be derived from anatomical scans processed with pipelines that are robust to scan quality (4,5). Retrospective analysis of clinical brain MRIs offers a complement to costly, prospective studies. This protocol outlines procedures to systematically, feasibly, and transparently screen clinical MRIs for inclusion into brain research using radiology reports.

Widescale implementation of retrospective studies that harness databases of clinical brain MRIs have the potential to augment existing pediatric brain MRI studies by several orders of magnitude. It is already commonplace to use hospital records for recruitment in brain imaging studies (6), but to date studies often stop short of incorporating brain MRIs archived in electronic health records (EHRs). For instance, The Children’s Hospital of Philadelphia (CHOP) and University of Pennsylvania previously drew upon CHOP clinical records for recruitment of the Philadelphia Neurodevelopmental Cohort (PNC) (7). The PNC included clinical phenotyping of approximately 10,000 participants of ages 8 to 21 – with neuroimaging in approximately 1,500 participants. Another widely used pediatric imaging cohort, the 21-site Adolescent Brain and Cognitive Development (ABCD) study, includes brain MRIs from approximately 11,000 participants (8). Although large by pediatric neuroimaging research standards, these sample sizes are dwarfed by the number of clinically-acquired brain MRIs contained within the EHR of a single hospital system (Figure 1).

The use of clinical scans also has the potential to increase the generalizability of neuroimaging findings beyond the demographically narrow population subset that typically participates in prospective neuroimaging studies. It has been well-documented that individuals who tend to participate in neuroscience and psychology studies are often demographically unrepresentative of clinical populations to whom the research is argued to generalize (9). Repurposing clinical neuroimaging may partially mitigate the disparity in resources affecting many families’ and individuals’ abilities to participate in research (Table 1). Moreover, the incorporation of clinical scans may improve the inclusion of lower functioning children in research, as well as individuals with neurogenetic conditions who are of particular interest for many researchers (10–12).

**Table 1.** Demographics of Available Clinical MRI Sessions.

| **Characteristic** | **Overall**<br>N = 232,474<sup>†</sup> | **Sex** |  |
| --- | --- | --- | --- |
|  |  | **Female**<br>N = 113,953<sup>†</sup> | **Male**<br>N = 118,521<sup>†</sup> |
| **Number of Individuals Scanned** | 127,643 | 64,852 | 62,791 |
| **Age at Imaging Session** |  |  |  |
| 0-5 years | 83,703 | 36,742 | 46,961 |
| 6-10 years | 50,543 | 22,850 | 27,693 |
| 11-15 years | 51,319 | 25,733 | 25,586 |
| 16-20 years | 30,667 | 16,791 | 13,876 |
| 21 years and older | 16,242 | 11,837 | 4,405 |
| **Race/Ethnicity** |  |  |  |
| American Indian or Alaska Native | 168 | 98 | 70 |
| Asian | 7,363 | 3,509 | 3,854 |
| Black or African American | 39,100 | 19,396 | 19,704 |
| Hispanic or Latino | 22,324 | 11,111 | 11,213 |
| Multiple Races | 5,024 | 2,410 | 2,614 |
| Native Hawaiian or Other Pacific Islander | 121 | 69 | 52 |
| Other | 13,725 | 6,433 | 7,292 |
| Unknown | 3,815 | 1,936 | 1,879 |
| White | 140,834 | 68,991 | 71,843 |
| <sup>†</sup> n |  |  |  |

Despite this promise, methodological procedures describing how to utilize clinical brain MRIs are limited. Many patients who receive clinical brain MRIs ultimately are found to lack any pathology that would exclude them from participating as a “healthy control” in a research study. “Clinical controls” may thus consist of patients who underwent brain MRIs to rule out serious neuropathology – for example, patients with headaches, possible head trauma, or developmental concerns – who were scanned and found to have unremarkable MRIs (13). Once a large dataset of clinical controls is identified, it can be harnessed as a comparison group for a specific population of interest. Identifying these clinical controls at scale, however, is hindered by the lack of a robust system to identify clinical brain MRIs without imaging pathology. In this protocol, we detail a process of identifying these “scans with limited imaging pathology” (SLIPs) through manual review of signed radiology reports in the EHR by study team members without formal clinical training using a predefined categorization scheme.

## Methods

In the present protocol, guidelines to grade radiology reports were developed through consultation with neuroradiologists at CHOP and Penn (JES, AV, SS), with iterative reviews and updates. A snapshot of our grading guidelines as of May, 2025, is publicly available (https://github.com/BGDlab/radiology_report_grading/blob/main/guidelines.pdf). Radiology report graders are given guidelines to grade each report as a 0, 1 or 2. In summary, a grade of 0 indicates that a report details serious imaging pathology or suggests an unusable scan due to image artifacts. Keywords or phrases indicative of gross pathology or permanent brain changes result in a grade of 0 (e.g. periventricular leukomalacia, volume loss, or glioblastoma; Figure 2). For convenience, severe motion artifacts or other notable problems with scan quality are also graded as a 0. Any language suggesting history of neurosurgical intervention (e.g. mention of craniectomy or ventriculoperitoneal shunt) is also grounds for a grade of 0. In contrast, a grade of 2 indicates there was no reason to suspect imaging pathology and that it is unlikely these patients would be excluded as a control sample in a research study (Figure 3). An absence of pathology indicated by phrases such as ‘unremarkable MRI of the brain’ or ‘normal anatomic variant’ results in a grade of 2. Common incidental findings such as pineal cysts or Virchow-Robin spaces and extracranial pathology do not preclude a report from receiving a grade of 2 (14–17).

In cases where it is not clear whether a report is 0 or 2, a rating of 1 is assigned (Figure 4). These reports may or may not be included as controls in downstream analyses depending on the goal of the analysis, the specific pathology noted, and any associated diagnoses. Observations such as benign enlargement of the subarachnoid space, grey matter heterotopia, or Chiari Type I malformation result in a grade of 1 as these findings may or may not differentiate these patients from typically developing children (18). Mention of microcephaly or macrocephaly in the radiology report is also grounds for a grade of 1 due to expected differences in brain volume (though extreme cephalic conditions such as those present in neurogenetic syndromes are graded as 0). Additionally, minor imaging artifacts from orthodontic hardware or mild patient motion result in a grade of 1 due to the potential of biases in downstream analysis (19,20). In the rare cases when a grader is unable to grade a scan due to ambiguity or complexity of overlapping pathologies/artifacts, the report is “skipped” and queued for team review. Weekly meetings are then held where skipped reports are reviewed and discussed until a consensus is reached and a grade can be assigned. In rare cases where a consensus cannot be reached, the case is escalated to a neuroradiologist for review. Any updates to the grading guidelines are added after the review and regrading.

Graders are instructed to consider all aspects of a report when assigning grades. In many hospital systems, radiology reports follow a semi-structured schema that typically includes six sections (though the exact labels may vary between radiologists): Indication, History, Comparison, Technique/Procedure, Findings, and Impression (21). The Clinical Indication contains information on why the MRI scan was ordered, such as suspected conditions (e.g., epilepsy) or reported problems (e.g., headaches, altered vision). This section may also contain pertinent medical history, including diagnoses from previous encounters. Comparisons include information on previous scans that can be used to monitor progression of pathology, such as the date of a previous MRI scan. The Technique includes information on the scanning sequences run, contrast administration, magnetic field strength of the system, and sometimes diagnostic limitations of the examination. Clinical brain MRIs typically include T1-weighted, T2-weighted, fluid attenuated inversion recovery (FLAIR), diffusion-weighted, and susceptibility-weighted images. At CHOP since 2008, routine 3 Tesla brain MRIs have typically included a standardized high-resolution, isotropic T1-weighted magnetization prepared rapid acquisition gradient echo (MPRAGE) sequence (22). The Findings and Impression sections make up the bulk of each report, with the Findings detailing the presence or absence of relevant pathologies appreciated on imaging given the clinical context and the Impression summarizing the derived conclusion from the findings.

Though no formal parsing of the report into sections is performed, most grading decisions arise from the content in the Findings and Impression sections. In some cases, graders also take into consideration aspects of the Technique and History when rating reports. When prior scans are mentioned in the History or Comparison sections along with phrases like “no new pathology noted”, graders should infer previous pathology not explicitly mentioned and grade the report as a 0 or 1. In practice, unusually long reports tend to indicate a degree of concern based on patient history or prior examinations. Thus, graders will assign a grade of 0 or 1. Technical aspects of a report also influence its grading as scans limited to specific anatomical regions like the orbits, auditory canals or mandibular regions are graded as 0 because they do not capture the whole brain. Certain protocols or sequences that do not result in a full high quality anatomical MRI data are also graded as a 0 (e.g. sessions only included a limited surgical navigation scan). Information on patient behavior leading to early scan termination or incomplete sequence runs due to an inability to tolerate results in a grade of 0 if it is clear from the report that there is no usable or inadequate anatomical scan. Disorders or diseases noted in the Indication section are only considered if they are pathognomonic of brain pathology. In general, the Indication is not used as the basis for grading a scan (e.g traumatic brain injury indication is not confirmation of a diagnosis if other sections do not mention it). Subtle findings in sequences not tailored for evaluation of grey and white matter (e.g., magnetic resonance angiography (MRA)) or variant anatomy vascular findings on MPRAGE are not considered when assigning a grade (see Figure 3B) (23,24). More concerning findings noted on these sequences however, such as an aneurysm or cerebrovascular occlusion, result in a grade of 0. Occasionally, reports note a need to communicate abnormal findings with patients as mandated in Pennsylvania by PA Act 112. Though not a diagnostic term, mention of this notification warrants caution, often leading to grade downgrades (i.e., grade of 1 downgraded to a 0).

### Grading Pipeline

The full pipeline for grading radiology reports is described in Figure 5. First, a SQL database of patients with MR imaging procedures is defined using EHR data stored in EPIC Clarity, containing radiology reports, demographics, and other clinical encounter data (25). The database is managed by Arcus, a suite of tools and services developed to enhance research efforts at The Children’s Hospital of Philadelphia. The data management framework includes 1) user access controls; 2) patient privacy and confidentiality protections through regulatory review; 3) electronic honest-brokered data de-identification and re-identification; and 4) data retention, management, sharing, and destruction services in an auditable computational environment. Data on Arcus is managed through the oversight of the CHOP Institutional Review Board, and access is governed by multiple institutional policies. Arcus security configuration and controls are based on the HIPAA Security Rule and are subject to audit by CHOP’s independent Internal Audit department, which reports directly to the CHOP Board of Trustees. Data is preserved according to international digital archiving standards, including replication across geographically disparate, secure, monitored storage environments to prevent loss in a catastrophic event. The virtual lab environment provided by Arcus provides access to a Linux based system with access to Jupyter, RStudio, and SQL.

The radiology report grading workflow then utilizes Python-based functions to create an interactive Jupyter notebook to present radiology reports and record assigned grades (https://github.com/BGDlab/radiology_report_grading) (Figure 5). First, procedure orders are filtered to identify radiology reports with anatomical MR imaging of the brain. Metabolic, functional, spectroscopy and post-mortem imaging are excluded. The reports associated with the filtered procedure orders are added to a SQL table that is accessed each time a grader initiates the grading process. Given the large quantity of reports available for grading, MRI procedures are then prioritized for specific projects based on patient age, sex, diagnosis, and other relevant criteria. For example, a project on preterm birth might prioritize neonatal scans where EHR information is also available on birth weight and gestational age at birth. The grading interface is accessed via a Jupyter notebook from within Arcus, in which the grader specifies the number of reports they want to grade in a batch and the project to pull those reports from. Presenting reports to grade in smaller batches prevents errors with reports loading when there is a long amount of time between a grade being entered and the reports being queued, which could occur when grading in large batches (n > 50) as each report can take between 1-3 minutes to grade. Data including the grade, the identity of the grader, the date of grading, and the project for which the scan was graded are stored in SQL tables on Arcus described above. A data management platform such as REDCap is an alternative to the Arcus platform that would be generalizable to other institutions.

Reports are presented for grading one report at a time, prioritized by the project-specific criteria such as year of procedure, with more recent scans appearing first. Highlighted text quickly directs the grader to keywords and phrases meaningful for determining a report’s grade (Figure 6). Strings that typically indicate section headers (e.g. Technique or Indication) are highlighted yellow, keywords indicating pathology or surgery such as ‘volume loss’ or ‘craniotomy’ in red, keywords indicating the absence of pathology such as ‘unremarkable’ and ‘within normal limits’ in green, and frequently used phrases that do not indicate pathology such as ‘ventricular system normal’ and ‘there is no mass effect or midline shift’ in gray. Graders are prompted to input a grade for each report twice to prevent errant entries. Recorded grades are added to a SQL table along with the grader’s name and datetime graded. When reports are skipped, indicated by a grade of −1, graders must list a reason why the report is being skipped.

Each report is assessed by at least two graders as a form of grade validation. SLIPs are identified as reports where two or more graders assigned the report a grade of 1 or 2. If either grader labeled the report as a 0, the report is excluded. Optionally, a natural language processing (NLP) model may be employed to act as one grader, with a human grader serving to validate the NLP grade, as described in Daniali et al (26). In brief, the NLP automates the classification of radiology reports, leveraging the detailed manual annotations. The pipeline begins with the fine-tuning of BERT-based language models (BERT, BioBERT, ClinicalBERT, and RadBERT) on a subset of our curated dataset with over 40,000 annotated pediatric radiology reports. The validation strategy accounted for class imbalance with a minority class weighting strategy, using train/validation/test splits and a cross-validation approach. Performance was evaluated and reported on each random seed and their average using accuracy, precision, recall (sensitivity), and F1-score. The pipeline showed a high degree of accuracy and robustness in accurately categorizing “normal” or “abnormal” reports across various experimental conditions, including out-of-distribution testing on reports from unseen reporting years, demographics, and hospital systems (25).

Scan sessions that meet criteria for SLIP are then requested from the radiology department with the aid of an honest broker: a third party who deidentifies DICOM metadata using Locutus (https://github.com/BGDlab/Locutus) (27) and plays no further role in downstream analyses. It is important to note that identifying “limited imaging pathology” is not necessarily indicative of a “clinical control”. For instance, depending on the research question, the appropriate reference group or “clinical control” can be further clarified based on International Classification of Disease (ICD) codes, clinical laboratory results, or review of physician notes. Additionally, manual or machine-assisted annotations of specific findings in radiology reports can be used as inclusion or exclusion criteria.

Data is eventually curated into a standardized format and then undergoes rigorous quality control before scans are processed and used for analysis. Raw imaging data is curated using HeuDiConv (28) and CuBIDS (29) with custom filters based on sequence acquisition parameters to robustly classify highly diverse clinical acquisitions into the standardized brain imaging data structure (BIDS) (30). Heterogeneous data is simplified by the identification of common sequences with similar imaging parameters (e.g. repetition time, voxel size, etc.) that are used to remove low-quality data. Scans with a low number of volumes - sometimes indicative of a prematurely terminated scan - are removed while scans with small parameter deviations (i.e. repetition of 3.002 seconds instead of 3 seconds) are preserved. The custom CuBIDS and HeuDiConv configurations used can be found here (https://github.com/BGDlab/chop-bgd-image-curation). Curated images are processed with FreeSurfer and SynthSeg+, a deep learning segmentation tool robust to clinical scan quality (4). A combination of manual and automated quality control can further refine SLIP cohorts. In a previous iteration of this protocol, images were manually rated by two independent raters to remove low-quality scans (22). Further, we employ the web application Swipes for Science (31) to scale manual review of large SLIP cohorts behind internal firewalls. Automated quality control utilizes the built-in SynthSeg+ (4) QC module to further refine SLIP cohorts. QC module and/or the Euler number from Freesurfer’s cortical reconstruction to further refine SLIP cohorts. To date, our focus is in using anatomical scans, either 1) focusing on a standardized T1-weighted MPRAGE sequence available in large subset of scan or 2) taking the median of all segmented volumes across all anatomical scans in a session (including T1-weighted, T2-weighted, and FLAIR images) as previously described (4). However, other brain MRI modalities are also available. For instance, standardized diffusion sequences are collected routinely at CHOP, which can be processed using robust pipelines including recently-developed and tractometry methods (32–34). Depending on the research question and downstream analyses, additional harmonization may be done using any one of the ComBat family of statistical harmonization tools to address scanner effects in derived phenotypes (35–40).

### Training Graders

Radiology report grading guidelines are designed to be used by non-expert graders with a range of educational backgrounds, including undergraduate students, graduate students, postdoctoral fellows, medical students, and non-radiologist physicians. The training process occurs in three distinct steps: introduction to radiology reports and review of grading guidelines, grading practice and self-evaluation, and an evaluation of reliability. Prospective graders meet with a senior grader to receive a formal introduction to the grading process. Once the introduction process is complete, prospective graders practice grading a predefined set of 50 radiology reports that are annotated with grades and grade justifications from previous graders to familiarize themselves with the grading process. Next, the prospective grader completes a self-evaluation, in which the grader completes 100 report grades and receives instant feedback after each grade through comparison with previous graders. This process allows the prospective grader to modify their grading strategies based on instantaneous feedback. Finally, there is an evaluation of grader reliability in which the prospective grader evaluates 150 reports without instantaneous feedback. Following this evaluation, Cohen’s kappa is calculated as a measure of inter-reliability between a new grader’s grades and those of senior graders for the reliability reports (41). A new grader’s reliability is confirmed once the kappa statistic is greater than 0.8 compared to the consensus among senior graders, indicating high inter-grader reliability (41). (Nine of the 150 reports have been flagged by senior graders as particularly challenging; these reports are reviewed with new graders but are excluded from the calculation of the kappa statistic.) In cases where reliability does not meet the threshold, new graders meet with a senior grader to review reports with major discrepancies (e.g. senior grader rated a report as 0 and new grader rate report as 2).

## Discussion

As of 2025, The Children’s Hospital of Philadelphia (CHOP) electronic health record (EHR) contained records of over 250,000 brain MRIs that have been qualitatively reviewed for clinical purposes and then archived. These pediatric MRIs can be repurposed for neurodevelopmental research, using the radiology report grading protocol detailed here to identify scans with limited imaging pathology (SLIPs). One future direction is for the resulting manual grades to serve as training data for natural language processing tools to automatically identify SLIPs, which can be applied to other hospital systems with minimal effort (26).

One application of SLIP data is to construct clinical brain growth charts. A precursor to the present protocol was previously used to derive clinical brain growth charts for 532 clinical controls aged 0-22 years (22,42). Encouragingly, clinical brain growth charts were convergent with growth charts derived from large-scale research datasets in terms of the normative developmental trajectories predicted by the models for global and regional brain phenotypes (22,42). Further, there was no evidence of age or sex biases relating to specific scanners or to manual or automated measures of image quality (22). The clinical indication of the scans did not significantly bias the output of clinical brain charts in preliminary data (22). Since this proof-of-concept publication, more than 10,000 additional SLIP data points have been identified using the present protocol. Scans of patients with specific diagnoses or genetic variants can now be benchmarked to these growth charts to quantify patterns of brain development in specific populations (43).

A limitation of the present protocol is that it relies on the content of radiology reports. While comprehensive, these reports are clinical summaries and might not always capture every subtle imaging feature that a direct image review by a neuroradiologist might reveal. An important validation will be to compare annotations of clinical summaries with direct image review in a subset of scans.

In summary, the present protocol provides a methodology to identify clinical brain MRI scans with limited imaging pathology (SLIPs) using EHR data. The protocol offers a promising avenue for quantitative neuroimaging research to augment prospective research datasets, leveraging the extensive repositories of scans routinely acquired by hospital systems in the course of clinical care (44). Harnessing clinical MRIs is a highly cost-efficient strategy to increase sample sizes and to expand generalizability to clinical populations. Ultimately, this protocol aims to enable scalable, reliable identification of clinical control brain MRIs, supporting large-scale, generalizable neuroimaging studies of typical brain development and neurogenetic conditions.

## Acknowledgements

This work was funded by the CHOP Research Institute and NIMH R01MH134896 (PI Alexander-Bloch). Additional support was provided by the Penn-CHOP Lifespan Brain Institute (LiBI) and NIMH R01MH120482. The authors acknowledge undergraduate research assistants who contributed to developing grading guidelines including: Madison Dengel, Harry Hearn, Naomi Shifman, Julia Katowitz, Alexa DeJean, Sayyid Khalil, Milo Writer, Olivia Scott, Vyvy Mai.

## Reproducibility

Code for radiology report grading and for image curation will be publicly shared upon publication at https://github.com/BGDlab/radiology_report_grading and https://github.com/BGDlab/chop-bgd-image-curation.

## Ethics and Dissemination

This protocol was reviewed and determined to be exempt from further oversight by the Children’s Hospital of Philadelphia’s Institutional Review Board. The exemption includes a waiver of informed consent as the study consists of retrospective data with low risk to human subjects, and requiring informed consent would bias the resulting sample of brain MRIs (IRB 20-017870). Datasets generated with this protocol cannot be made available for public access as they contain protected health information from CHOP patients.

---

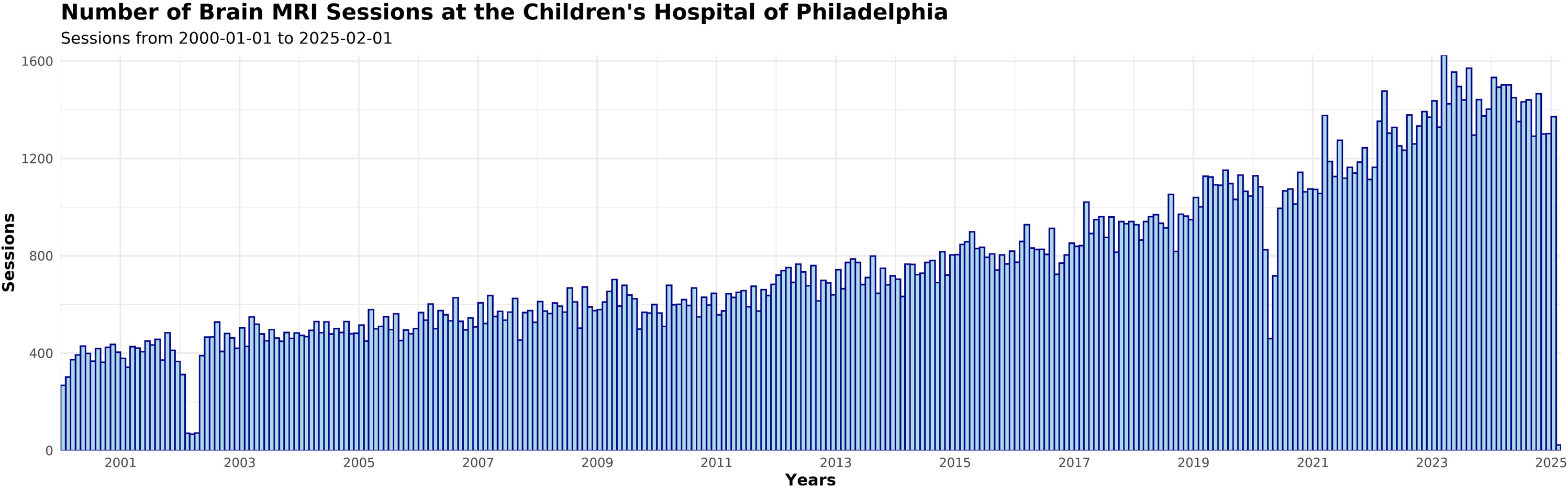

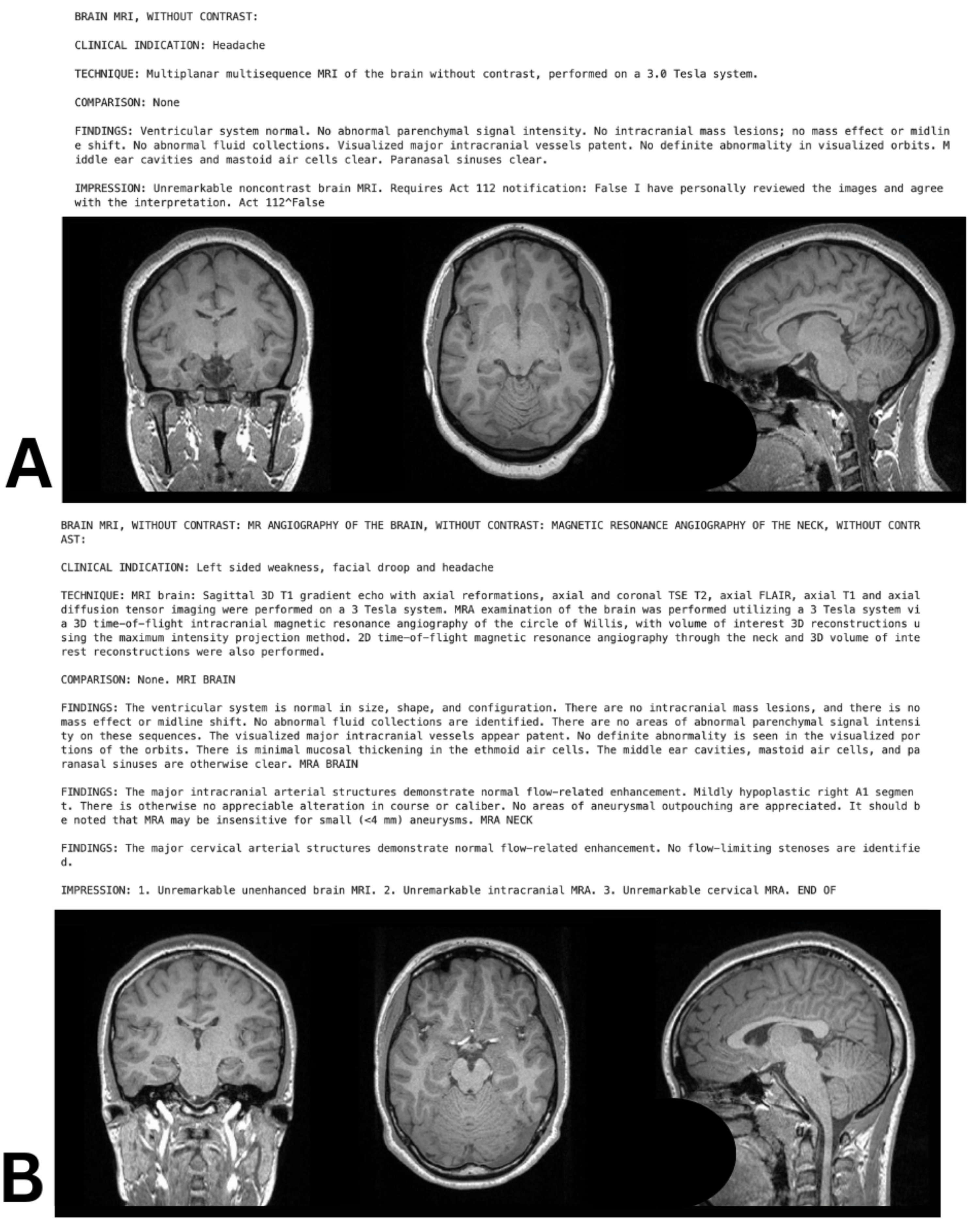

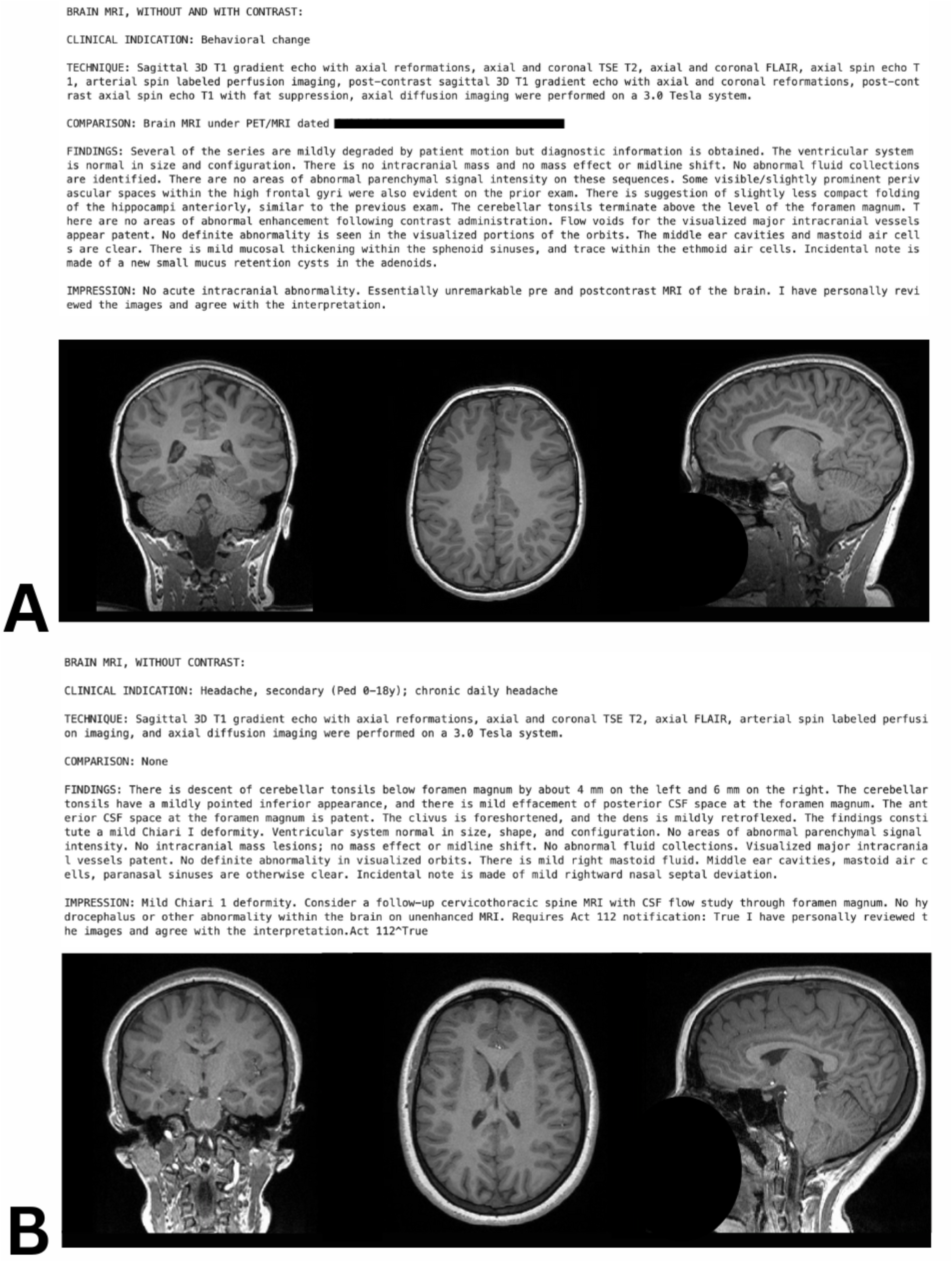

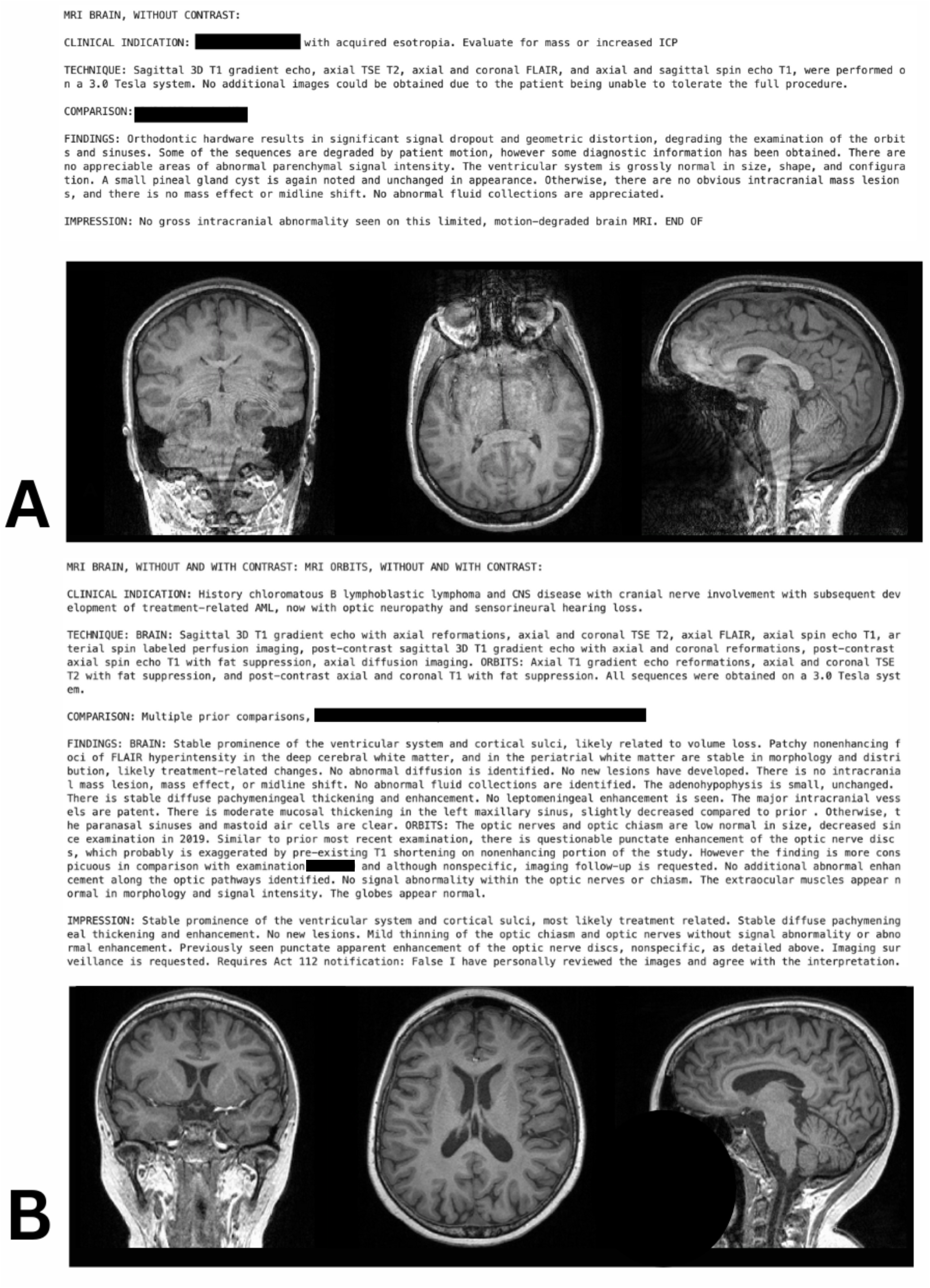

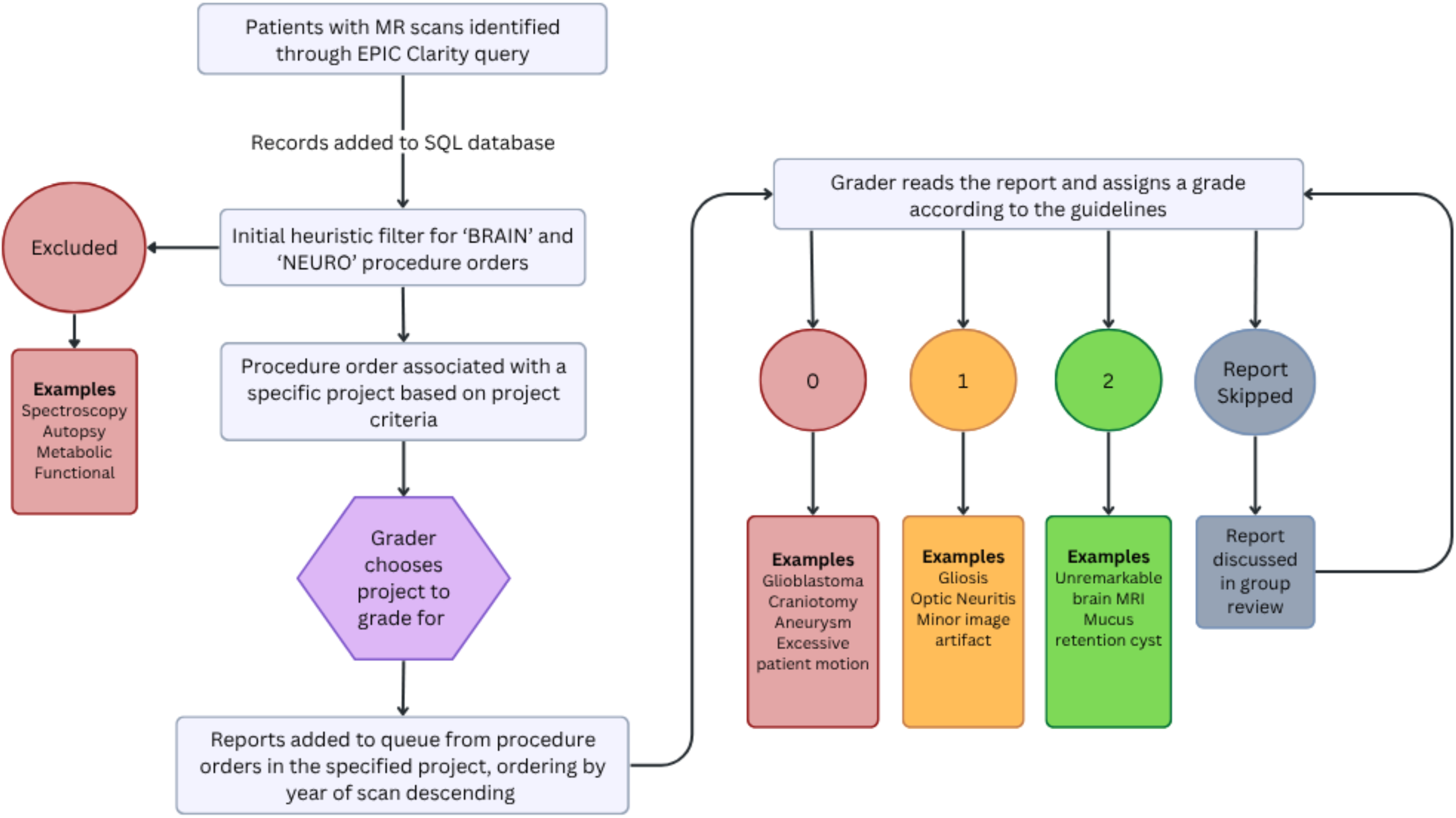

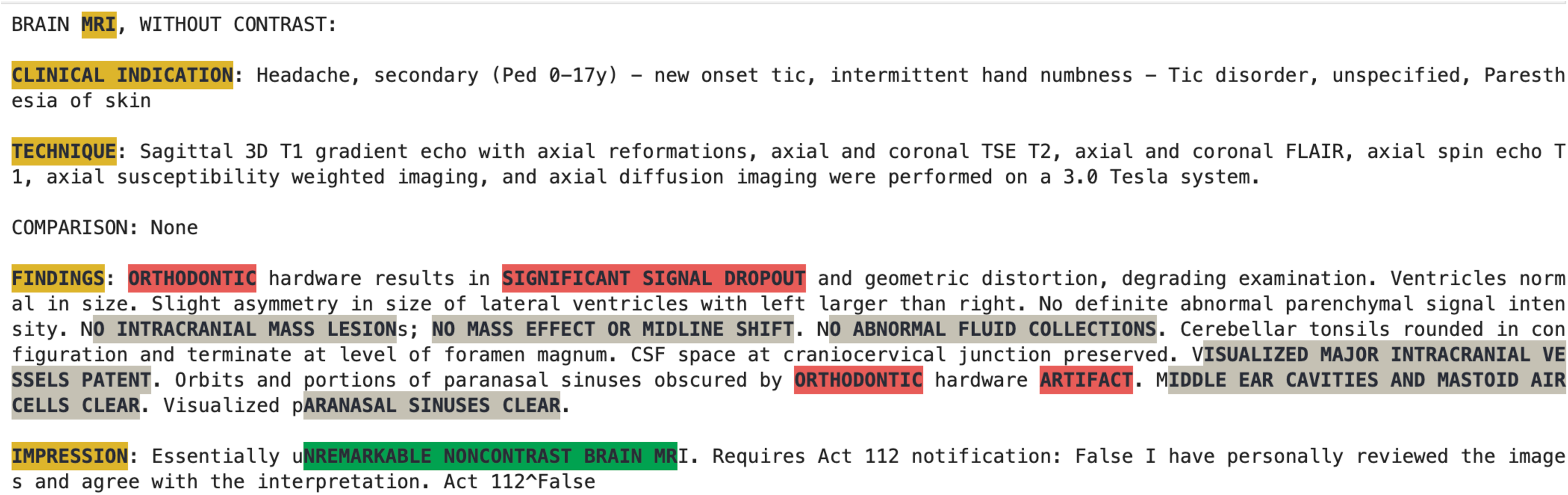

